# A genome-annotated bacterial collection of the barley rhizosphere microbiota

**DOI:** 10.1101/2021.03.10.434690

**Authors:** Senga Robertson-Albertyn, Federico Concas, Lynn H Brown, Jamie N Orr, James C Abbott, Timothy S George, Davide Bulgarelli

## Abstract

We generated a bacterial collection from the rhizosphere of cultivated barley (*Hordeum vulgare* L. ssp. *vulgare*) to assess taxonomic distribution of culturable members of the barley microbiota and their plant growth-promoting potential. From this we retrieved strains belonging to the dominant phyla of the plant microbiota— Actinobacteria, Bacteroidetes, Firmicutes and Proteobacteria—and gathered evidence they code for functional genes implicated in nitrogen fixation, hydrogen cyanide channels and phosphate solubilisation. Here we present an initial comparative genomic analysis of the collection revealing that plant growth-promoting potential of the culturable barley bacterial microbiota appears to have a relatively broad phylogenetic base while retaining some strain-specificity.

## MAIN TEXT

The microbial communities thriving at the root-soil interface, collectively referred to as the rhizosphere microbiota, represent an untapped resource of plant probiotic functions [1, 2]. Bacterial members of the microbiota capable of enhancing plant’s mineral uptake from soil and pathogen protection, collectively referred to as plant growth-promoting rhizobacteria (PGPRs) have gained prominence in both basic science and translational applications [3-5].

As a resource for comparative investigations of the plant microbiota across host species, we provide a collection of 53 bacterial strains, isolated from the rhizosphere of barley (*Hordeum vulgare* L. ssp. *vulgare*), the fourth most cultivated cereal worldwide [6]. Bacteria were isolated from the rhizosphere of plants grown in an agricultural soil previously used for investigations in barley-microbiota interactions [7, 8]. To facilitate further experimentations, we subjected individual isolates to whole-genome sequencing and we performed a preliminary comparative analysis which identified varying presence of genes associated with PGPRs.

Using the table of taxonomic ID to Kyoto Encyclopaedia of Genes and Genomes (KEGG) ortholog (KO) identifier, we predict 6,429 unique KOs across all isolates. When examining the KOs that overlapped between isolates it was observed that there was a group of 514 KOs that were conserved between all isolates (Figure 1). In the case of Microbacterium and Stenotrophomonas groups, all isolates assigned to each group shared the same KOs based on this analysis. However, this same observation did not extend to isolates assigned to the Pseudomonas and Bacillus groups: Bacillus samples had 240 gene intersections shared only between themselves. In addition, among the analysed *Pseudomonas* sp, where 4 isolates share 24 KOs that did not feature in any other sample (within or outwith *Pseudomonas* sp.), which were assigned to 3 different *P. brassicacearum* isolates according to 16S rRNA gene identity (bi75, bi06, bi11, bi55). A further *P. brassicacearum* isolate, bi70, assigned to yet another OTU, had its own set of 24 unique KEGG orthologs versus the other isolates.

**Figure 1.**
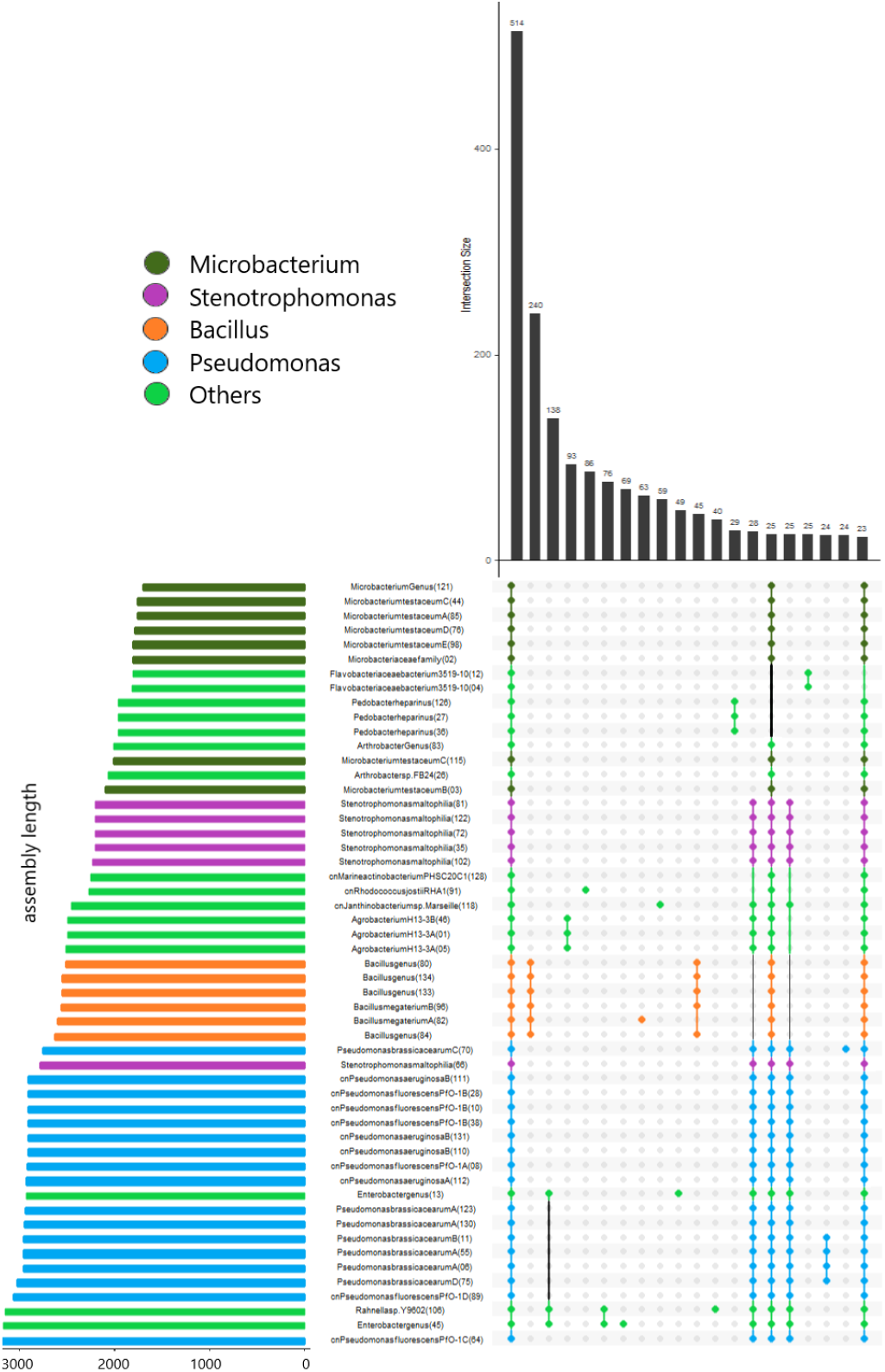
KEGG ortholog intersection map of all high-quality genome isolates. Colours depict the assigned genus of each isolate based on Amphora taxonomic assignment from full genome sequence reads. Intersection size denotes the number of KEGG orthologs shared by the isolates indicated by the corresponding circle below the intersection. The number in brackets next to the isolate description denotes the isolate ID and the length of the coloured bar next to each isolate represents the length of the sequence assembly of the individual sample.

In order to identify bacterial isolates that may have downstream applications e.g. in inoculant trials, we compared whole-genome average nucleotide identity (ANI) across isolates which allowed us to identify individual “founder” members (ANI cut-off 96%, Figure 2). Next, we further inspected these genomes, representing 11 distinct genera, for the presence of known PGPR genes focussing on, *dinG, hcn, nif, pho* and *pqq* (Table 1). These genes have been previously implicated in the acquisition of the essential elements phosphorus and nitrogen and enhancing the efficacy of biological fertilisers to enhance plant nutrient uptake [9-13]. Congruent with the overall KO analysis, we detected within genera variation for the plant-growth promoting potential of the identified isolates. For instance, our results showed that a variety of *pqq* genes belonging to the PQQ operon were present in a number of bacterial isolates. Notably, *ppq*B, C, D, E, F and (in 7 isolates) H were almost exclusively found together in *Pseudomonas* derived isolates, apart from the bi13, Enterobacterium genus, isolate in which *ppq*B, C, D, E and F were also detected. Although the majority of isolates did not carry any *ppq* gene orthologs, in addition to these there were 4 isolates that carried a single *pqq* gene ortholog (Arthrobacter FB24 (bi26); *pqq*F, *Stenotrophomonas maltophilia* (bi66); *pqq*F, *Pseudomonas aeruginosa* (bi112); *pqq*B and *Janthinobacterium* (bi118); *pqq*L and one (*Microbacterium testaceum* (bi03)) that carried a pair of *pqq* gene orthologs, *pqq*C and *pqq*D. Interestingly, *pqq*E was not detected in this isolate when it has been demonstrated that *pqq* genes are present in a conserved order and that, occasionally, *pqq*CDE can be fused with *pqq*D being fused to the C-terminus of *pqq*C and/ or the N-terminus of *pqq*E [14]. Likewise, we observed between genera variation for genes putatively implicated in nitrogen metabolism. For instance, isolate bi46 *Agrobacterium* H13-3 was identified as carrying the most *nif* gene orthologs, *nif*ALSU. However, 20 isolates across a variety of genera including *Janthinobacter, Pedobacter* and *Pseudomonas* were observed to have *nif*A orthologs which have previously been identified in and identified in nitrogen-fixing *Pseudomonas* [15] as well as in bacteria implicated in denitrification [16].

**Figure 2.**
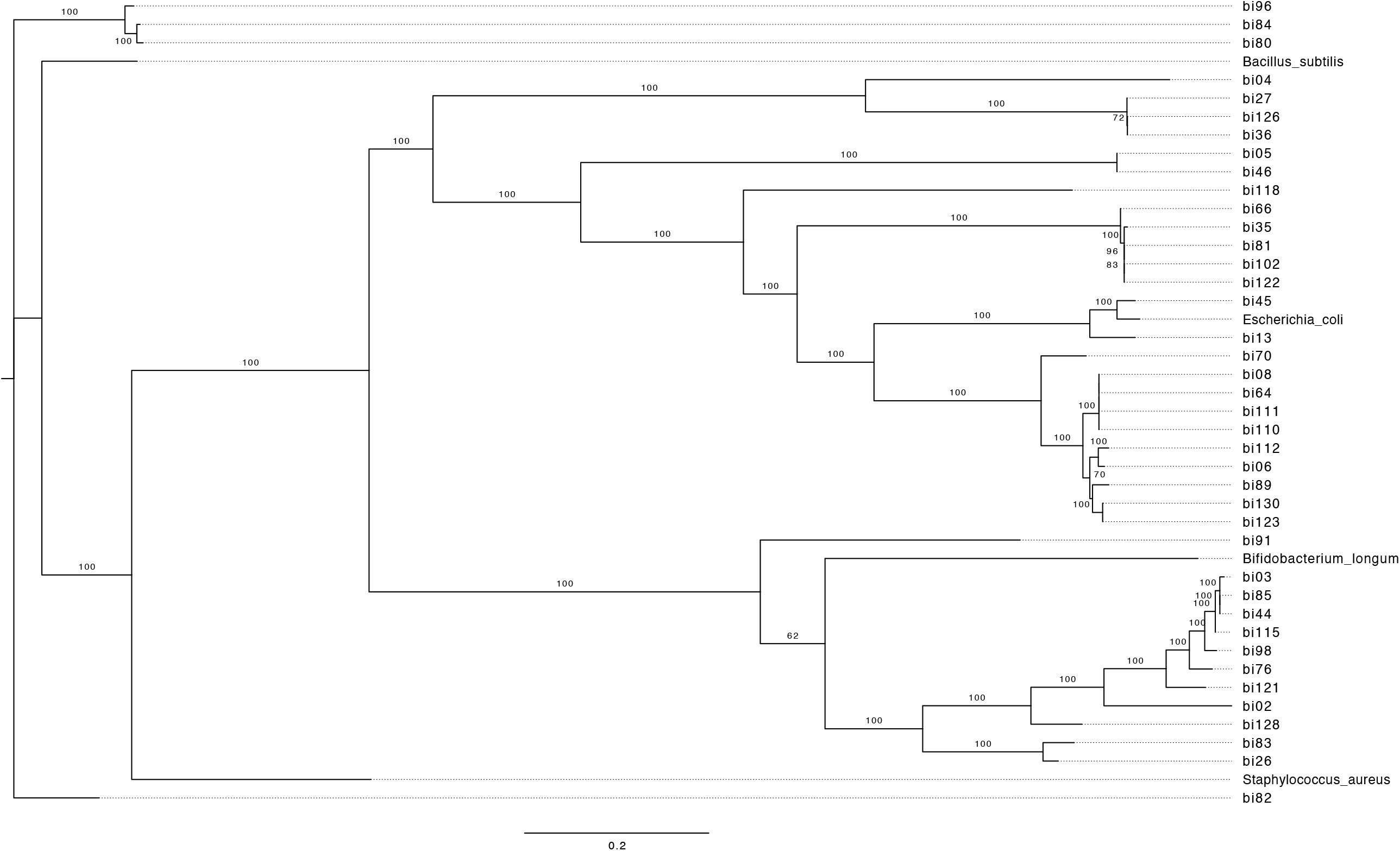
Whole genome phylogenetic tree of individual bacterial genomes (ANI cut-off 96%) constructed incorporating additional sequences for *Bacillus subtilis* NCIB 3610 (GCA_006088795), *Bifidobacterium longum* NCC2705 (GCA_000007525), *Escherichia coli* K12 MG1655 (GCA_000005845) and *Staphylococcus aureus* NCTC 8325 (GCA_000013425). Protein predictions were obtained using Prokka 1.14.6 and the tree was constructed with 100 bootstrap iterations.

**Table 1.**
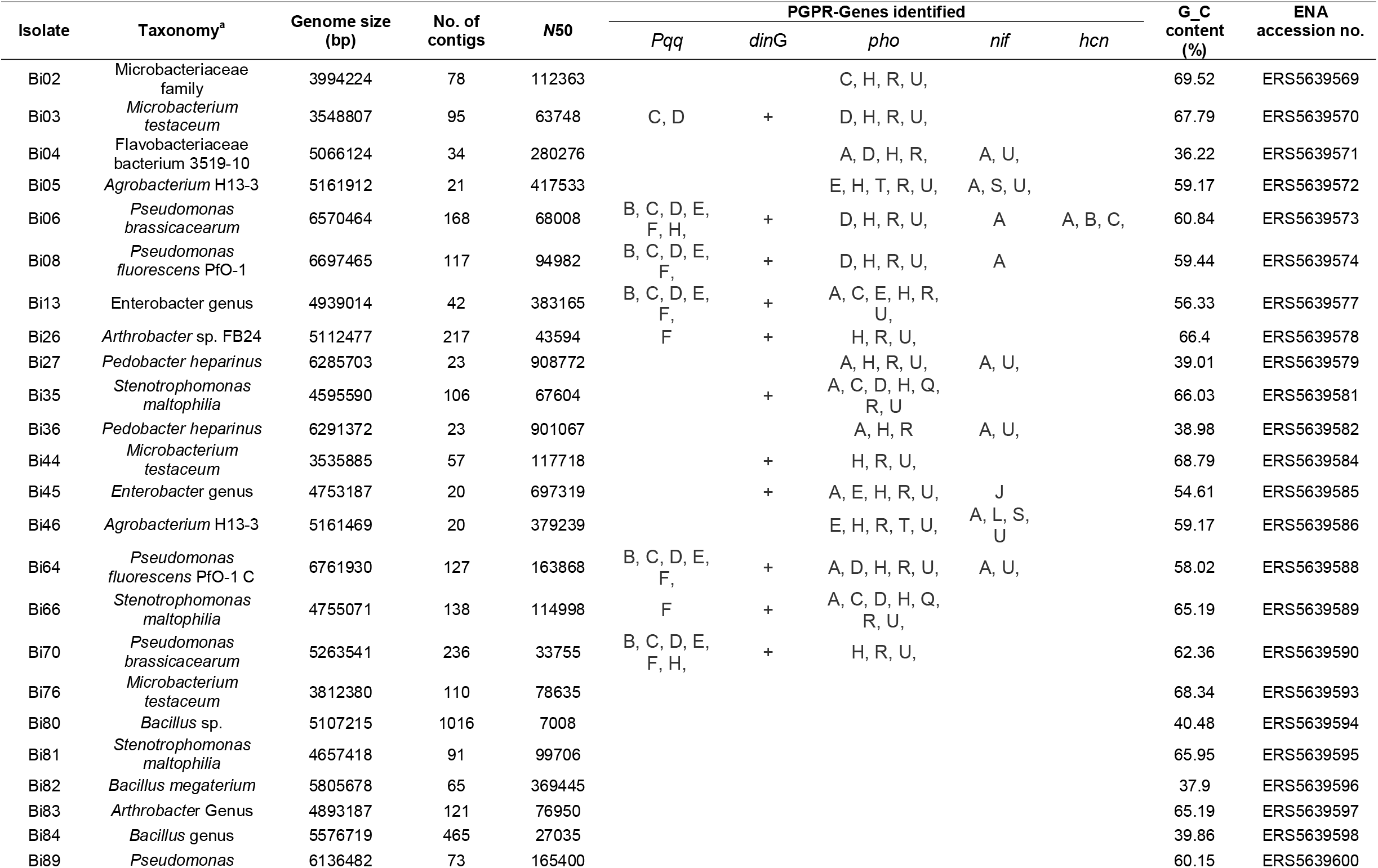
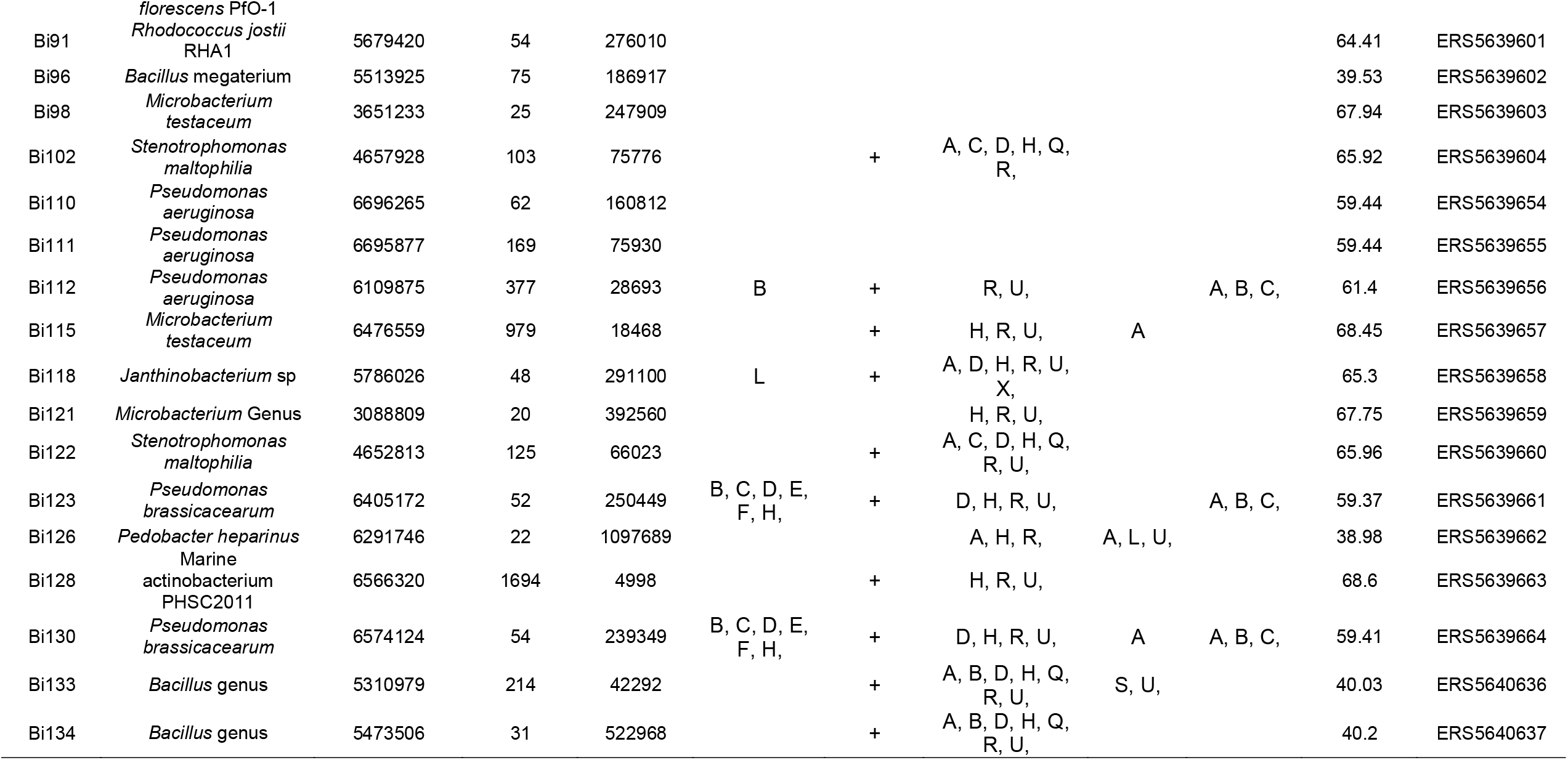
Taxonomic affiliation, genomic characteristics and accession numbers of genomes of 41 individual bacterial isolates (ANI cut-off 96%) described in this study. Capital letters depict actual genes identified within the inspected metabolic processes. For the *dinG* gene, the identification in each bacterial genome is depicted by the plus sign. ^*a*^Strain taxonomy reflects the lowest and unique rank representing over 90% of the amphora classification.

Taken together, our data indicate that the growth promoting potential of the barley bacterial microbiota appears to have a relatively broad phylogenetic base while retaining some strain-specificity, a similar observation has been recently reported for the root microbiota of the model plant *Arabidopsis thaliana* [17]. Our collection complements Synthetic Communities (SynComs) already available for the root endospheric microbiota of important cereal crops, such as maize [18] and rice [19], and it is ideally placed to foster further collaborative studies in molecular plant-microbe interactions.

## METHODS

Rhizosphere specimens were prepared as in previous studies [7, 8] and serial dilutions of rhizosphere suspension were plated on Nutrient [20] or R2A [21] agar plates and incubated for 48 -72 hours at either 27 °C or at room temperature (approximately 20 °C). We initially selected 194 distinct CFUs on the basis of different morphology and colour. Of these, we retained 84 strains for full genome sequencing. Individual bacterial genomes were sequenced using the ‘standard service’ of MicrobesNG (Birmingham, UK; https://microbesng.com/). Briefly, bacterial genomic DNA libraries were prepared using Nextera XT Library Prep Kit (Illumina, San Diego, USA) following the manufacturer’s protocol with the following modifications: 2 ng of DNA instead of 1 ng were used as input, and PCR elongation time was increased to 1 min from 30 seconds. DNA quantification and library preparation were carried out on a Hamilton Microlab STAR automated liquid handling system. Pooled libraries were quantified using the Kapa Biosystems Library Quantification Kit for Illumina on a Roche light cycler 96 qPCR machine. Libraries were sequenced on the Illumina HiSeq using a 250bp paired end protocol. Reads were adapter trimmed using Trimmomatic 0.30 with a sliding window quality cut off of Q15 [22]. De novo assembly was performed on samples using SPAdes version 3.7 [23], and contigs were annotated using Prokka 1.11 [24]. On the basis of GC content and unambiguous taxonomic annotation generated using amphora [25] classification, we retained 53 genomes for downstream analyses. To compare only components of characterised metabolic pathways, predicted genes from all isolates were concatenated and annotated with eggNOG-mapper (v1.0.3) [26]. The resultant annotation file was parsed in Python to generate a table of taxonomic ID to Kyoto Encyclopaedia of Genes and Genomes (KEGG) ortholog (KO) identifier. From this table a presence-absence matrix of all KOs predicted at least once in each isolate was generated in R. To compare all predicted proteins between isolates, predicted proteomes were clustered using OrthoFinder (v2.2.1) [27], and functionally annotated using InterProScan (v5.29-68.0) [28]. Clusters and annotations were aggregated using KinFin (v1.0) [29]. Cluster and KO intersections were plotted using UpSetR (v1.3.3) [30]. The phylogenetic tree was constructed using bcgTree 1.1.0 [31] and RAxML 8.2.12 [32], using RAxML’s GTRGAMMA model and 100 bootstrap iterations.

## DATA AVAILABILITY

The genomic sequences reported in this study are deposited in the European Nucleotide Archive (ENA) under the study number PRJEB42773. Accession numbers for the individual genomes of the “founder” members are provided in Table 1.

## AUTHORS CONRTIBUTION

SRA, TSG, DB: study design. SRA, FC, LHB: bacterial collection construction. SRA, JNO, JCA: genome data analysis. SRA and DB wrote the manuscript with inputs from all co-authors.

## ACKNOWLEDGEMENTS

We are thankful to Federica Caradonia (University of Modena and Reggio Emilia, Italy) and Carmen Escudero-Martinez (University of Dundee, UK) for their technical assistance during the development of the collection. This work was supported by a BBSRC iCASE studentship awarded to DB (BB/M016811/1) and partnered by the James Hutton Limited (Invergowrie, UK). FC was supported by an Erasmus+ Traineeship programme (European Commission). LB was supported by a James Black Prize Studentships (University of Dundee). JNO was supported by an ERC advanced grant “Shuffle” (Project ID: 669182) awarded to Robbie Waugh/The James Hutton Institute. SR-A, JA and DB are currently supported by the Horizon 2020 Framework Programme Innovation Action ‘CIRCLES’ (European Commission, Grant agreement 818290) awarded to the University of Dundee. James Hutton researchers receive financial support from the Rural and Environment Science and Analytical Service Division of the Scottish Government.

